# Random Mutagenesis for the Generation of Repertoire of Aureochrome-based Optogenetic Scaffolds

**DOI:** 10.64898/2025.12.26.696586

**Authors:** Madhurima Khamaru, Pracheta Pal, Saptarshi Pal, Devrani Mitra

## Abstract

Engineering of the natural light-regulated transcription factors for the purposeful generation of synthetic photo-switches is a widely adopted strategy in synthetic biology and optogenetics. Aureochrome is one such potential, naturally occurring photoreceptor-cum-transcription factor that can be used for the development of synthetic optogenetic tools, especially meant for light-regulated gene expression. It is however well-known that different biological events demand different duration of signaling state lifetime and diverse affinity towards different DNA substrates. Therefore, in this study, we produce multiple variants of aureochromes via random mutagenesis - mimicking single generation directed evolution. After screening the single generation variants for in-frame transcripts, we sort them on the basis of altered photocycle kinetics as well as DNA-binding affinity. The variants that exhibited reversible photochemistry and light-regulated DNA-binding ability similar/comparable to that of the wild-type despite incurring mutations, are characterized and discussed in this manuscript. These can later be subjected to successive rounds of random mutations to get an eventual superior variety with enhanced functionality. Even the apparently unsuccessful ones, which depicted drastic alteration of photochemical/DNA-binding properties, helped us to identify amino acid residues – lesser known to be indispensable for a certain biological activity. The process of diversification and selection via random mutagenesis not only explains the functional significance of the different amino acids in aureochromes but also generates a repertoire of appropriate aureochrome variants that may be used in optogenetics.

## 1 Introduction

Directed evolution brings the essence of natural evolution into the laboratory (Beaudry and Joyce, 1992) and remarkably accelerates the origin of new variants, which might take thousands of years in nature (Scientific Background on the Nobel Prize in Chemistry, 2018) (Kauffman, 1993; Wells, 1990). This strategy of protein engineering is helpful in the selection of a candidate with the most desirable traits, particularly when there is no previous information on the structural and functional details of the wild-type protein. The desired variant can be obtained through iterations of diversification followed by extensive screening and selection. The workflow for diversifying the starting nucleotide sequence involves two major approaches: error-prone PCR (McCullum et al., 2010) and gene shuffling (Stemmer, 1994). While error-prone PCR is undoubtedly an efficacious way of incorporating random changes in an existing protein sequence, gene shuffling enables wider chunks of similar genes to be swapped, considerably expanding the repertoire of variations produced (Cobb et al., 2013). Error-prone polymerase chain reaction (epPCR) techniques are adaptations to standard PCR protocols that are intended to elevate the inherent error rate of the polymerase (Cirino et al., 2003). Error-prone PCR exploits the innate low fidelity of *Taq* DNA polymerase, which can be reduced further by adding manganese chloride (MnCl2) and/or an excess of magnesium chloride (MgCl_2_), and providing uneven amounts of the four dNTPs in the PCR mixture (Wilson and Keefe, 2001). For this study, error-prone PCR using MnCl_2_ and/or increased concentration of MgCl_2_ was employed to facilitate the incorporation of random point mutations in the gene of interest. Studies have demonstrated that substituting Mn^2+^ for Mg^2+^ has a negative impact on both the fidelity of polymerase activity and the proofreading specificity of DNA polymerase I (El-Deiry et al., 1984). However, manganese probably does not hamper nucleotide selection because, when used in minute concentrations, Mn^2+^ enhances the fidelity of DNA polymerase I. At physiologically relevant concentrations (< 100 µM), manganese seems to introduce errors, supposedly by means of interaction with the DNA template. Further mutagenesis ensues at higher concentrations of Mn^2+^ (500 µM and 1.5 mM), possibly stemming from interactions of the ion with single-stranded regions of the DNA or certain sites of DNA polymerase I (Beckman et al., 1985). When used in optimal concentrations, Mg^2+^ serves as a cofactor of DNA polymerase, thereby boosting its enzymatic activity (Masoodi et al., 2021). During amplification, the magnesium ion binds to the alpha phosphate of a dNTP, facilitating the elimination of beta and gamma phosphates. The resultant dNMP forms a phosphodiester linkage with the 3’ OH group of the neighboring nucleotide. MgCl_2_ also plays a crucial role in facilitating the annealing of primers to their complementary regions in the DNA template. Mg^2+^ lowers the electrostatic repulsion between the DNA strands by associating with the negatively charged phosphate of DNA. Nevertheless, an excess of MgCl_2_ in the PCR mixture leads to non-specific annealing of primers, thereby triggering errors in DNA amplification (Excedr, 2022).

Aureochromes are unique photoreceptor-cum-transcription factors exclusively found in marine photosynthetic heterokont algae (Takahashi et al., 2007). They exhibit the rarest association of a light-oxygen-voltage (LOV) sensor domain for blue light (BL) reception (Mitra et al., 2012) with a basic-leucine-zipper (bZIP) motif meant for binding to specific DNA substrates (Matiiv and Chekunova, 2018). The distinctive co-occurrence of the LOV and bZIP domains, not known to exist in any photoreceptor apart from aureochromes, makes them BL-regulated transcription factors (Kroth et al., 2017). The LOV sensor in aureochrome undergoes a photochemical reaction upon BL absorption that transmits a signal via linker to its effector bZIP for DNA binding (Banerjee et al., 2016). Thus, it can act as an excellent photoswitch for attaining a quick change in gene expression by readily binding to DNA possessing the ACGT core sequence in response to BL without undergoing an elaborate signaling cascade. Since aureochrome can be genetically encoded, placing the transgene under a native promoter in a target cell can bring about tissue-specific gene regulation, and orthogonal manipulation can drive non-invasive control over regulatory networks (Khamaru et al., 2025). The photosensitivity of LOV and the lifespan of the photoproduct are vital for the proper functioning of aureochromes. Further, its light-dependent DNA-binding efficiency is the most crucial factor to consider for its use as an optogenetic tool. Optogenetics (Emiliani et al., 2022) amalgamates genetics with optics to facilitate precise, non-invasive, spatiotemporal, and reversible regulation of gene expression in response to light. Naturally occurring photoreceptors or light-sensitive proteins and their engineered counterparts are instrumental in the development of optogenetic tools (Zhao et al., 2025). Optogenetics relies on design and engineering of existing proteins (Banerjee and Mitra, 2020) and nature proves to be the greatest designer at any time. Over the years nature continues to innovate, evolve and optimize novel proteins with improved functionalities. In fact, all the natural proteins are the products of successive natural mutations followed by the natural selection of the beneficial mutants. Evolution keeps on generating mutations. Though not all the sequences encode functional proteins, evolution continues with the selection of those limited numbers of meaningful functional proteins. Directed evolution mimicking this process of natural evolution brings a new fitness landscape in terms of functionality through the imposed artificial selection (Arnold, 2012). Therefore, an error-prone PCR based technique of random mutagenesis (Leung et al., 1989) has been adopted here being inspired by the approach of ‘directed evolution’ for the development of an appropriate optogenetic scaffold from the BL-responsive photoreceptor, aureochrome (Aureo). Random mutations were incorporated into the wild-type aureochrome gene sequence by epPCR to create a repository of diverse aureochrome sequences, each coding for a different mutated version of the same protein. The mutants that hold the potential to express functionally active proteins were then screened and grown in culture to determine their functional characteristics. It is therefore fascinating to witness how a single generation of random mutagenesis alters the fitness landscape of Aureos with desired/improved photochemical and DNA-binding properties. The eventuality of these induced changes determines the hotspots present in the linker, LOV, and bZIP domains of aureochrome and reflects the importance of those amino acids in its functionality. This, in turn, paves the way for the use of this algal photoreceptor-cum-transcription factor as an optogenetic scaffold.

## 2. Materials & Methods

### 2.1. Error-prone polymerase chain reaction (epPCR)

The synthetic gene corresponding to the coding region of wild-type aureochrome of alga *Thalassiosira oceanica* (*To*AubZL), containing the bZIP and LOV domains of *To*Aureo along with the linker, was taken as the template for PCR amplification. In order to introduce random point mutations into the wild-type gene, different concentrations of manganese chloride (MnCl_2_) (SRL) and/or an excess of magnesium chloride (MgCl_2_) (SRL) were added to the error-prone PCR mixture.

### 2.2. Cloning into expression vector pET28a

The epPCR products were purified and then subjected to restriction digestion by BamHI and SacI enzymes (New England BioLabs). Every double-digested DNA fragment was ligated to the pET28a vector at BamH1 and Sac1 restriction sites. These ligated constructs were then transformed into the DH5α strain of *Escherichia coli* competent cells. Correctly cloned products were selected by performing colony PCR and were validated through sequencing.

### 2.3. Sequence analysis of the generated variants

DNA sequence chromatogram files from Applied Biosystems DNA sequencers were analyzed using Chromas 2.6.6 (https://technelysium.com.au/wp/chromas/). The nucleotide sequences were translated into protein sequences using the ExPASy Translate tool (https://web.expasy.org/translate/). Protein sequence with the desired open reading frame (ORF), i.e., starting with methionine, ending at a stop codon, and containing polyhistidine-tag (6xHis-tag) at the N-terminal, was submitted to BLASTX (https://blast.ncbi.nlm.nih.gov/Blast.cgi?PROGRAM=blastx) as query. Sequences with no mutation and mutated sequences with premature stop codons leading to truncated proteins were rejected. Multiple sequence alignment of all the variants with the wild-type aureochrome protein was carried out on the Clustal Omega server (https://www.ebi.ac.uk/Tools/msa/clustalo/). The positions of occurrence of the random mutations were documented.

### 2.4. Protein overexpression and purification

The mutated DNA constructs were transformed into the C43 strain of *E. coli* for protein expression. Colonies obtained from the transformation plate were inoculated into 10 mL autoclaved Luria-Bertani (LB) broth culture containing 50 mg/mL kanamycin (Hi Media) and grown overnight at 37 ⁰C and 150 rpm in the temperature-controlled incubator-shaker (Thermo Fisher Scientific). The overnight culture was then transferred into 1 L of autoclaved LB broth containing 50 mg/mL kanamycin. Bacterial growth continued at 37 ⁰C and 160 rpm for ≈1.5 hours, until OD_600nm_ reached 0.6. Protein overexpression was then induced by adding 300 µM isopropylthio-β-galactoside (IPTG) (SRL). Following IPTG induction, the cells were grown in the dark at 22 ⁰C and 120 rpm overnight.

The induced culture was pelleted down in multiple rounds of centrifugation at 8,000 rpm for 10 minutes, at 4 ⁰C. The cell pellet was dissolved in a lysis buffer (20 mM Tris, pH 8; 50 mM NaCl; 10% glycerol), protease inhibitor cocktail (Sigma-Aldrich), a pinch of lysozyme (SRL), and a pinch of phenyl methyl sulfonyl fluoride (PMSF) (SRL) were added to the dissolved pellet prior to incubation on ice for ≈15–20 minutes. Sonication (Hielscher Ultrasonics) was performed on ice. The lysed pellet was centrifuged (Thermo Fisher Scientific) at 14,000 rpm for 1 hour, at 4⁰C. After centrifugation, the supernatant was subjected to nickel-nitrilotriacetic acid (Ni-NTA) affinity chromatography (Qiagen). After passing wash buffer (20 mM Tris, pH 8; 50 mM NaCl; 10% glycerol; 10 mM imidazole) through the Ni-NTA column, the protein fractions were collected using elution buffer (20 mM Tris, pH 8; 50 mM NaCl; 10% glycerol; 200 mM imidazole). The eluted fractions containing the protein of interest were passed through a sephadex desalting column for the removal of imidazole. The desalted fraction was concentrated using the Amicon centrifugal filter unit (Merck-Millipore). The purified protein samples were subjected to 15% sodium dodecyl sulfate-polyacrylamide gel electrophoresis (SDS-PAGE) and resolved at 120–150 V against a protein molecular weight marker (Bio-Rad).

### 2.5. Photochemical kinetics study

The absorption spectrum of each of the purified and desalted proteins was recorded using a UV-vis double beam spectrophotometer (Shimadzu, UV-2600) for the range 230-750 nm. After recording the absorption spectrum in the dark state, the protein was exposed to LED light for 10 minutes. Upon light absorption, aureochrome is known to undergo a conformational change associated with a formation of covalent adduct, leading to the disappearance of the triplet absorbance peak observed previously in the non-photobleached condition. Absorbance of the aureochrome protein in every consecutive cycle was recorded over a certain period of time, until the protein reverted back to its native state and regained the absorption maxima at 447 nm. For each *To*AubZL mutant, the duration of each scan was noted to calculate the dark recovery time of the protein from the value of its absorbance at 447 nm. The rate constant (*k*) for each construct was then calculated by plotting the absorbance value (OD_447nm_) against time using a single exponential equation [OD_447nm_ = constant(1-e^-*kt*^)], followed by fitting the graph to a mono-exponential decay function.

### 2.6. Estimation of protein concentration

Absorbance of the aureochrome protein at 447 nm was recorded using a UV-vis spectrophotometer (Shimadzu). Protein concentrations were determined by applying Beer-Lambert law, and extinction coefficient of 12,500 M^−1^ cm^−1^.

**A = ε c 1**

[A = Absorbance at 447 nm; ɛ = Extinction coefficient = 12,500 M^−1^cm^−1^; c = Concentration of the protein; l = Path length = 1 cm]

### 2.7. Electrophoretic mobility shift assay (EMSA)

Electrophoretic mobility shift assays (EMSAs) were performed for the quantitative assessment of protein-DNA interaction. Lyophilized single-stranded complementary oligonucleotides purchased from IDT/Eurofins were annealed in annealing buffer (10 mM Tris, pH 8.0; 20 mM NaCl). Protein-DNA interaction studies were performed with a final concentration of 0.5 µM double-stranded DNA in binding buffer (50 mM Tris, pH 8.0; 50 mM NaCl; 1.25 mM MgCl_2_; 0.01 mg/mL BSA; 20% glycerol). Serially diluted protein was added to the mixture of double-stranded DNA and binding buffer, followed by incubation on ice for 45 minutes under blue light exposure or in complete darkness. The free DNA as a control and protein-DNA samples in decreasing order of protein concentrations were loaded into separate wells of the 10% native polyacrylamide gel, and resolved at 130 V for 1 hour in 0.5X Tris-Borate-EDTA (TBE) buffer with 2 mM MgCl_2_. Bio-Rad Mini-PROTEAN Tetra Vertical Electrophoresis Cell and PowerPac Basic Power Supply were used for the electrophoresis, and the EMSA was performed at an ambient temperature of 4 ⁰C. After the completion of the electrophoretic run, the gel was stained using SYBR Gold (Thermo Fisher Scientific) in 0.5X Tris-Borate-EDTA (TBE) buffer, by placing it on a rocking platform for 45 minutes. The gel was imaged using a gel documentation system (Bio-Rad). K_D_ values were calculated using ImageJ and KaleidaGraph. The three-parameter Hill function equation i.e., Y = m1*x^m2 /(m3^m2 + x^m2) was used, where m1 represents the maximum specific binding, m2 is the Hill slope, and m3 is the K_D_.

## 3. Results

### 3.1. Identification and analysis of the mutants generated through epPCR

Four rounds of epPCR using different concentrations of MnCl_2_ and/or MgCl_2,_ introduced random mutations into the wild-type aureochrome gene (*To*AubZL) (**SI-1**)as evident from the screening and sequencing of the purified PCR products cloned into pET28a vector. The detailed domain-wise documentation of the mutations is represented in **Table 1**. Although various mutations were introduced across the LOV and bZIP domains, the S314C mutation in the bZIP domain emerged as the most frequent, appearing in three variants.

**Table 1:**
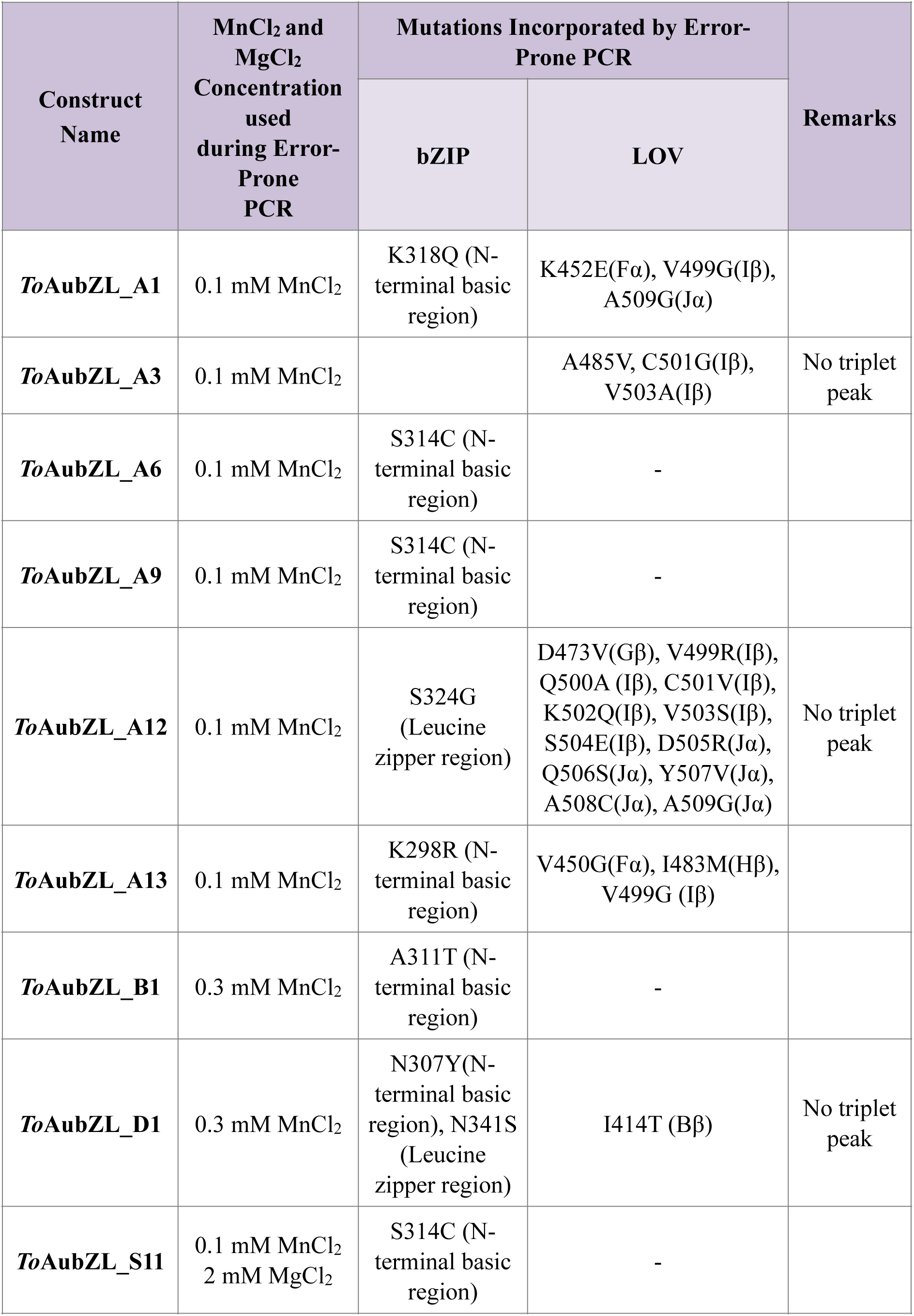
Summary of mutations found in the different domains of *To*AubZL variants.

| Construct Name | MnCl<sub>2</sub> and MgCl<sub>2</sub> Concentration used during Error-Prone PCR | Mutations Incorporated by Error-Prone PCR |  | Remarks |
| --- | --- | --- | --- | --- |
|  |  | bZIP | LOV |  |
| ***ToAubZL_A1*** | 0.1 mM MnCl<sub>2</sub> | K318Q (N-terminal basic region) | K452E(Fα), V499G(Iβ), A509G(Jα) | |
| ***ToAubZL_A3*** | 0.1 mM MnCl<sub>2</sub> | | A485V, C501G(Iβ), V503A(Iβ) | No triplet peak |
| ***ToAubZL_A6*** | 0.1 mM MnCl<sub>2</sub> | S314C (N-terminal basic region) | - |  |
| ***ToAubZL_A9*** | 0.1 mM MnCl<sub>2</sub> | S314C (N-terminal basic region) | - |  |
| ***ToAubZL_A12*** | 0.1 mM MnCl<sub>2</sub> | S324G (Leucine zipper region) | D473V(Gβ), V499R(Iβ), Q500A (Iβ), C501V(Iβ), K502Q(Iβ), V503S(Iβ), S504E(Iβ), D505R(Jα), Q506S(Jα), Y507V(Jα), A508C(Jα), A509G(Jα) | No triplet peak |
| ***ToAubZL_A13*** | 0.1 mM MnCl<sub>2</sub> | K298R (N-terminal basic region) | V450G(Fα), I483M(Hβ), V499G (Iβ) | |
| ***ToAubZL_B1*** | 0.3 mM MnCl<sub>2</sub> | A311T (N-terminal basic region) | - |  |
| ***ToAubZL_D1*** | 0.3 mM MnCl<sub>2</sub> | N307Y(N-terminal basic region), N341S (Leucine zipper region) | I414T (Bβ) | No triplet peak |
| ***ToAubZL_S11*** | 0.1 mM MnCl<sub>2</sub><br>2 mM MgCl<sub>2</sub> | S314C (N-terminal basic region) | - |  |

The variant *To*AubZL_A12 exhibited the highest number of mutations, the majority of which occurred in the LOV domain, including the Q500A mutation in the FMN-binding pocket of the LOV sensor. Two distinct positions within the LOV domain – Val499 and Val503 – can be regarded as mutational hotspots, as they were found to be mutated to different residues across various constructs. Val499 is mutated to arginine in *To*AubZL_A12 and to glycine in both *To*AubZL_A1, and *To*AubZL_A13; while Val503 is substituted by alanine in *To*AubZL_A3 and by serine in *To*AubZL_A12. No mutations were introduced in the LOV sensor domain of the variants *To*AubZL_A6, *To*AubZL_A9, *To*AubZL_B1, and *To*AubZL_S11. Some of the mutations were found in the FMN-binding pockets, while others were located in close proximity to the FMN-binding residues (**Figure 1a**). In some variants, mutations even occurred at residues that are otherwise conserved (**Figure 1a**).

**Figure 1:**
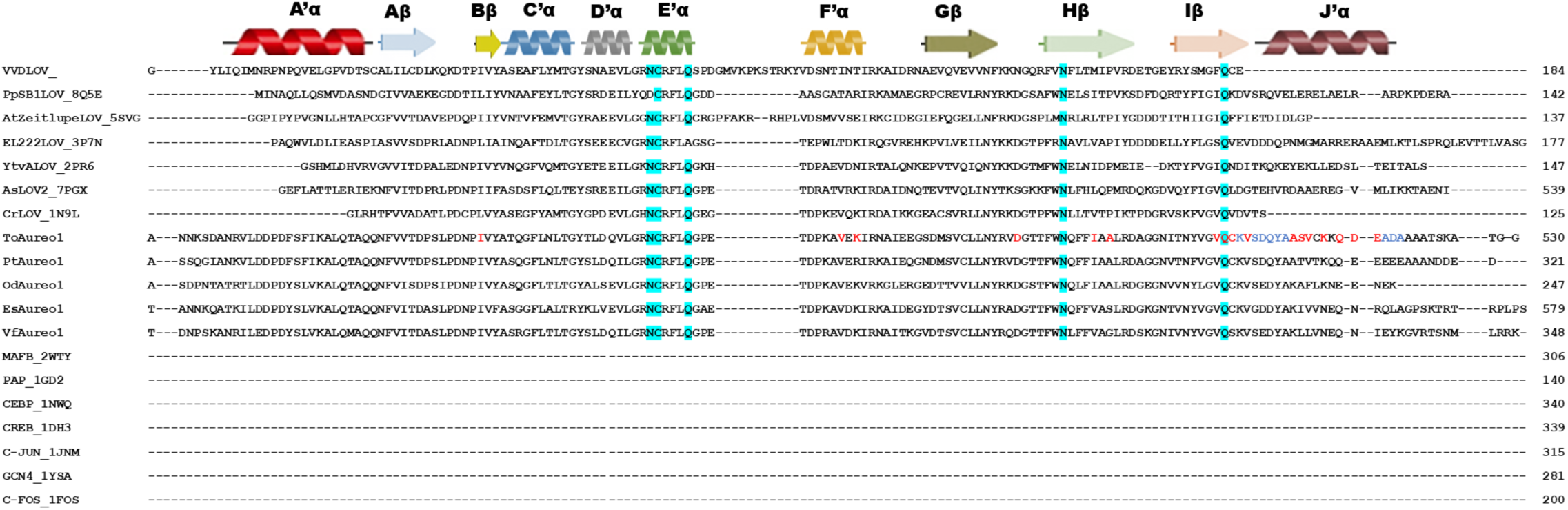
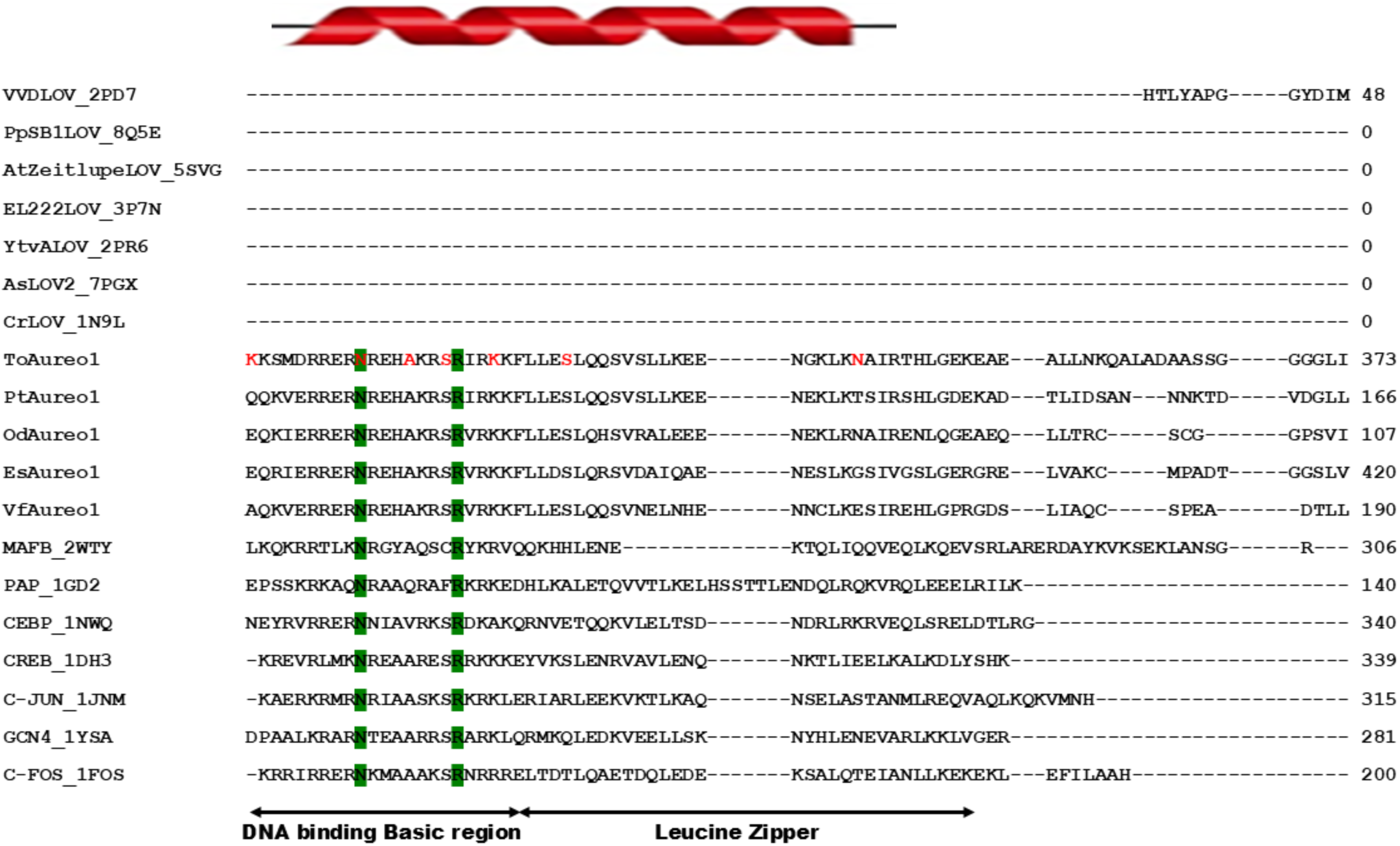
a. Multiple Sequence Alignment of the LOV domains from the *To*AubZL variants with other aureochromes and LOV proteins with available crystal structures in PDB to determine the residue differences. Residues of *To*AubZL that were found to be mutated in the variants generated through random mutagenesis are colored in red/blue. FMN-binding residues of the LOV domain are colored in cyan. b. Multiple Sequence Alignment of the bZIP domains from the *To*AubZL variants with other aureochromes and bZIP proteins with available crystal structures in PDB to determine the residue differences. Residues present at the boundary of the DNA-binding signature motif of bZIP are colored in green.

Mutations in the bZIP domain were identified in the majority of the variants, with the exception of *To*AubZL_A3. Some of the variants even had mutations in critical regions, such as the conserved signature motif of the DNA-binding basic region and the leucine heptad repeat, that is essential for the dimer stabilization (**Figure 1b**). *To*AubZL_A1, *To*AubZL_B1, *To*AubZL_D1, *To*AubZL_A6, *To*AubZL_A9, and *To*AubZL_S11 contain mutations in the signature motif of the DNA-binding basic region of the bZIP domain. The variants *To*AubZL_A6, *To*AubZL_A9 and *To*AubZL_S11 all carried a Ser314 mutation. This position is naturally occupied by cysteine in bZIP sub-groups Maf, JUN, and FOS (**Figure 1b**). *To*AubZL_D1 exhibited the Asp341 mutation in the dimerization domain of bZIP. In *To*AubZL_A12, a mutation Ser324 was identified in the bZIP domain, positioned in between the DNA-binding basic region and the dimerization domain. In *To*AubZL_A13, Lys298 mutation was found at a position in the bZIP domain that was not directly involved in the interaction with DNA. Among all the mutations introduced in the bZIP domain, N307Y found in *To*AubZL_D1 was perhaps the most significant, as asparagine was found to be conserved across any bZIPs (**Figure 1b**). In the variant *To*AubZL_B1, the mutation Ala311 was found in the signature motif of the DNA-binding basic region, where residue substitutions were found naturally in bZIPs, such as PAP and CEBP (**Figure 1b**).

Furthermore, phylogenetic tree constructed separately taking the LOV and the bZIP domains of all the *To*AubZL variants with other aureochromes, LOV and bZIP proteins reflected the impact of these random mutations in the phylogeny of *To*AubZL (**SI-2**, **SI-3**).

### 3.2. Photochemical study and determination of the dark recovery kinetics

The LOV photoreceptors exhibit an absorption spectrum with an absorption maximum at ∼447 nm in the dark state or ground state due to the presence of an oxidized flavin cofactor. Upon photoexcitation, a covalent linkage is formed between the C(4a) carbon atom of the flavin cofactor and the thiol moiety of the active site cysteine (Conrad et al., 2014). Formation of this cysteinyl adduct within the LOV domain results in a conformational change of the protein, leading to the disappearance of the triplet peak at 447 nm. Following the withdrawal of light source, the protein reverts back to its native state with the restoration of the absorption maxima in course of time. LOV domains thus exhibit reversible photochemistry (Chang et al., 2017). The deprotonation of the FMN-N5 atom and the thermal breaking of the flavin-cysteinyl covalent linkage leads to recovery of the dark state. The rate of reversion to dark state determines the photocycle kinetics. Although adduct formation occurs within a few nanoseconds to microseconds, the lifetime of the photocycle can take seconds to days depending on the protein. The photocycle kinetics is crucial for any photoswitch as it determines the duration of the signaling state or steady state (Hemmer et al., 2023). Light-responsive proteins like phototropin, EL222, PHY-B of *Arabidopsis* are designated as fast cycling as their light state lifetimes are in the order of seconds. ZTL, VVD, YtvA show slow cycles as their light state lifetime ranges from hours to days (Losi et al., 2003; Pudasaini et al., 2017; Zoltowski et al., 2007). Both the short- or long-signaling states are desirable in different optogenetic applications, depending on the context and the specific effector function being targeted. Very recently, the incredible kinetic diversity of LOV photoreceptors ranging from picoseconds to days, has been mapped on an evolutionary timescale spanning hundreds of millions of years (Herzog et al., 2025).

Here, we explored the signaling state lifetime by determining the rate constant of dark recovery in all the functional constructs. Each aureochrome variant was checked for the presence of the characteristic triplet absorption peak of LOV at 447 nm in the dark using UV-vis spectrophotometry (**Figure 2**). The variants *To*AubZL_A3, *To*AubZL_A12, and *To*AubZL_D1 failed to show a prominent triplet absorption peak with the absorption maxima at 447 nm (**Figure 2**). While *To*AubZL_A3 and *To*AubZL_A12 contain multiple mutations in the LOV sensor, the LOV domain of *To*AubZL_D1 has only a single mutation, I414T.

**Figure 2:**
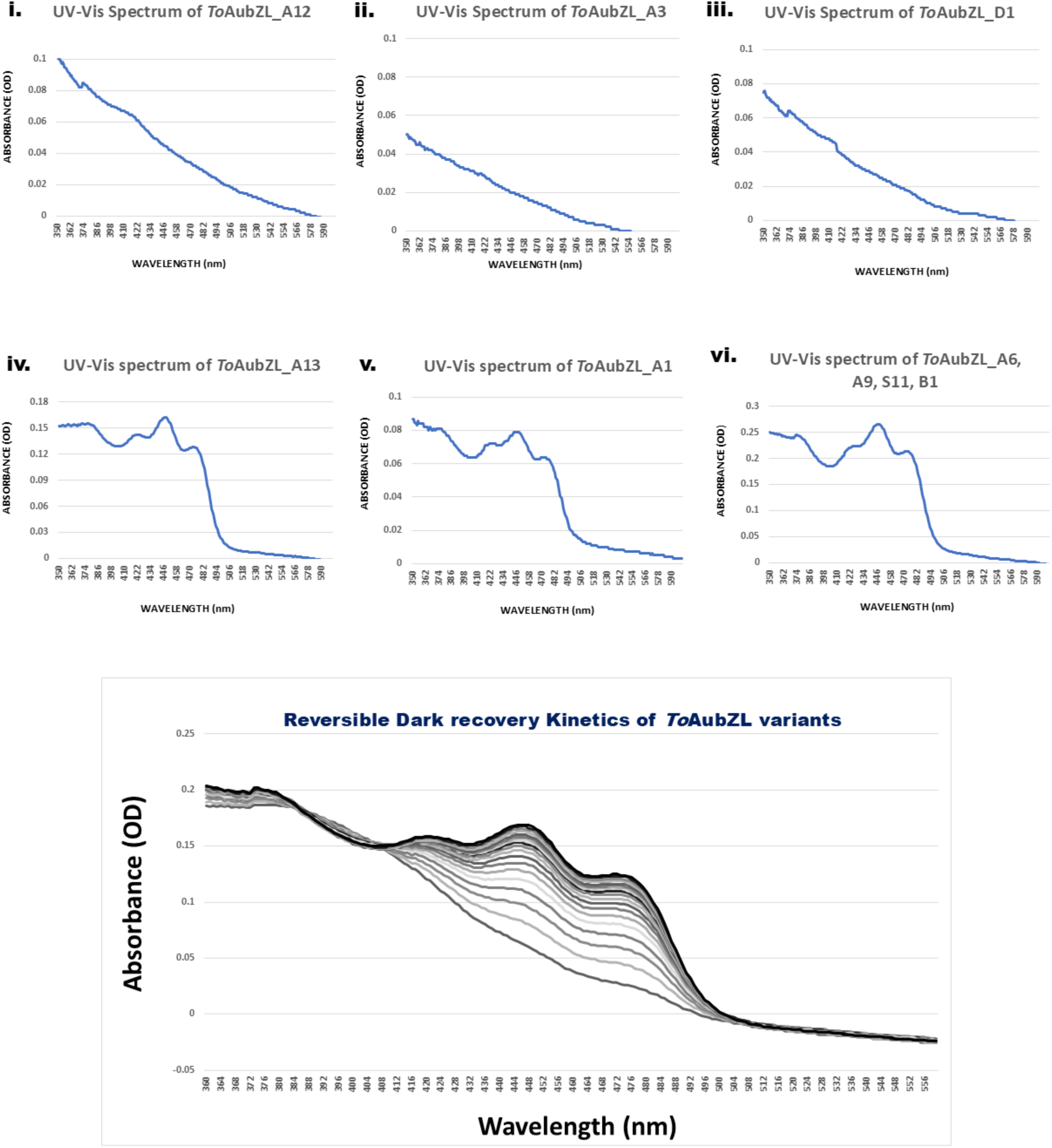
UV-Vis spectra of the LOV sensor of the variants of *To*AubZL.

In *To*AubZL_D1, Ile414 present at the vicinity of the FMN-binding pocket maintains hydrophobic contacts with the neighboring Val432 and Phe439, hence is essential (**Figure 3**). Incidentally, Phe439 is a part of the fully conserved GXNCRF motif, which is crucial for housing the flavin chromophore. In all LOV photoreceptors, the positions homologous to Ile414 are consistently occupied by either leucine or isoleucine. Similarly, Val432 position is mostly conserved for hydrophobic amino acid residues in the LOV photoreceptors. The mutation I414T significantly weakens the hydrophobic interactions, as shown in (**Figure 3**), while also enabling Thr414 to form an additional hydrogen bonding interaction with the sulfhydryl group of active site Cys437. Presence of a strong hydrophobic interaction network along with structured hydrogen bonding network is essential not only for the stability of the flavin chromophore, but also for effective signal transmission in LOV photoreceptors (Nakajima et al., 2021).

**Figure 3:**
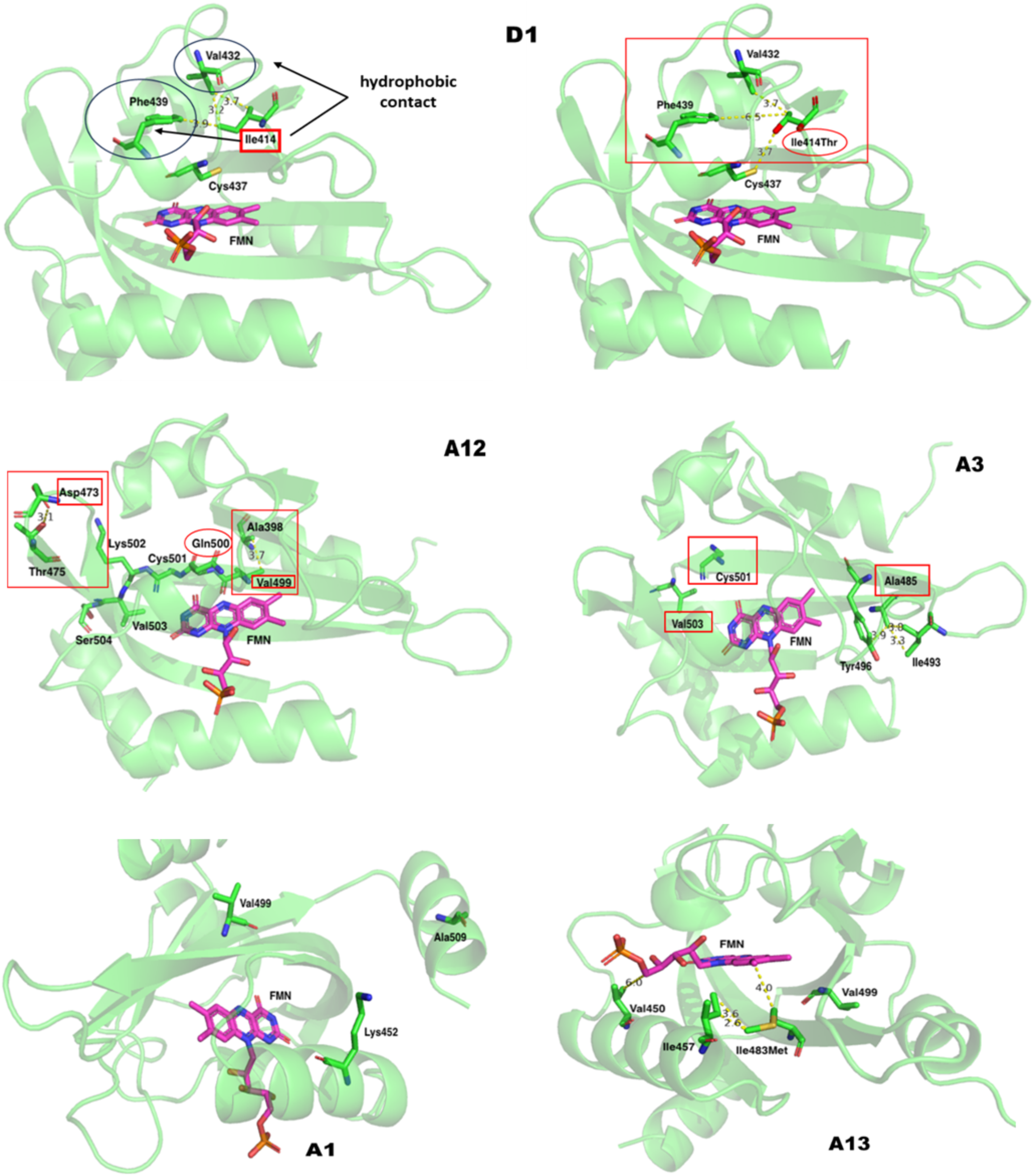
Structural insights into the LOV sensor of the *To*AubZL variants.

In *To*AubZL_A3, the mutations A485V, C501G, and V503A led to the loss of the characteristic triplet absorption peak of LOV around 447 nm. As reported for *Vf*Aureo1 (Nakajima et al., 2021) any substitution at the hydrophobic beta-sheet region can significantly alter the photochemical behavior of LOV photoreceptors. Incidentally, all these residues belong to the LOV core region. While Ala485 is located at Hβ, Cys501 and Val503 are located at the distal end of Iβ and the loop leading to Jα respectively. The length of the non-polar side chain of Ala485 is ideally suited to interact with Ile493 and the non-polar part of Tyr496’s phenyl ring (**Figure 3**), which in turn maintains hydrophobic contact with the isoalloxazine ring of FMN. Mutation to Val can possibly lead to steric clash with Tyr496, potentially disrupting the proper positioning of the flavin within the hydrophobic cleft. The specific effects of the mutations Cys501 to Gly and Val503 to Ala can be precisely understood upon availability of the experimental structure of *To*Aureo1. Aureo1. However, we speculate that exposure of the polar side chain of Cys501 to solvent and burial of the Val503 side chain impart the necessary conformational specificity at the Iβ–Jα region.

The variant *To*AubZL_A12, contains several mutations in the LOV domain – D473V, V499R, Q500A, C501V, K502Q, V503S, and S504E. While most mutations are located in the loop preceding Jα helix, two mutations – D473V and V499R – appear to be particularly critical. As already discussed, mutations within the LOV core, especially in the beta-core, can significantly alter flavin-binding and photocycle kinetics. The presence of the polar side chain of Asp473 (or its equivalent) is essential for maintaining polar contacts (hydrogen bonding) with the highly conserved Thr475 (or its polar/charged substitute at the homologous position) side chain as well as backbone (**Figure 3**). This interaction is critical for preserving structural stability in the loop region connecting Gβ–Iβ. Further, polar substituent at Asp473 position is necessary as the loop is solvent-exposed. Non-polar substitution like Val in this position is certainly unfavorable. Similarly, substitution of Val499 with Arg at Iβ significantly alters the former’s interaction with Ala398 near A’α. As amply suggested, hydrophobic interface is essential for LOV dimerization and signaling. Furthermore, in Aureos, the signal should propagate via A’α considering the presence of the effector bZIP at the N-terminus. Therefore, undoubtedly, Val499 substitution with a bulky and charged amino acid is likely to have serious consequences on the overall structural stability of the LOV core. However, when Val499 is substituted with another non-polar residue like Gly no serious effect is observed. We therefore speculate that these two mutations in combination provide fair justification for the absence of the characteristic triplet peak in the dark state. Mutants that failed to show absorbance maxima at 447 nm were excluded from further analysis.

The variants that exhibited the triplet absorption peak despite containing mutations in the LOV domain are *To*AubZL_A1 and *To*AubZL_A13 (**Figure 2**). The rate constant values (**SI-4**) are documented in **Table 2**. Variants with no mutations in the LOV domain exhibited dark recovery kinetics identical to that of the wild-type aureochrome. Interestingly, both *To*AubZL_A1 and *To*AubZL_A13 showed photocycle reaction kinetics at the order of 10^-3^, which is comparable to wild-type. Despite the mutations in the LOV domain, including residues involved in FMN-binding, these variants retained their photochemical properties, with only slight alterations observed in the rate constant of the photochemical cycle.

**Table 2:** Rate constants of the photocycle kinetics of different *To*AubZL variants.

| Construct Name | Rate Constant (S<sup>-1</sup>) |
| --- | --- |
| *ToAubZL_A1* | $2.15 \times 10^{-3} \pm 1.64 \times 10^{-4}$ |
| *ToAubZL_A3* | Triplet peak not found |
| *ToAubZL_A6* | $3.19 \times 10^{-3} \pm 1.04 \times 10^{-4}$ |
| *ToAubZL_A9* | $3.19 \times 10^{-3} \pm 1.04 \times 10^{-4}$ |
| *ToAubZL_A12* | Triplet peak not found |
| *ToAubZL_A13* | $1.26 \times 10^{-3} \pm 2.88 \times 10^{-5}$ |
| *ToAubZL_B1* | $3.19 \times 10^{-3} \pm 1.04 \times 10^{-4}$ |
| *ToAubZL_D1* | Triplet peak not found |
| *ToAubZL_S11* | $3.19 \times 10^{-3} \pm 1.04 \times 10^{-4}$ |

The mutations seen in the variant *To*AubZL_A1 are K452E, V499G, and A509G. The mutation V499G is common across several variants and appears to have no detrimental effect on the wild-type characteristics. K452E and A509G mutations have occurred at the two flanking ends of the LOV core, i.e., within the A’α and Jα helices, respectively (**Figure 3**). The homologous position of Lys452 is usually occupied by polar charged or even neutral residues in all LOV photoreceptors. In Zeitlupe, a glutamate is present naturally. Nevertheless, the long side chain of Lys452 is exposed to the solvent and buried towards the beta-core. Therefore, substituting Lys452 with another polar, charged or even neutral amino acid, like Asn or Thr, as found in some LOV domains, should not ideally impact the wild-type characteristics. Similarly, substitution of the buried Ala509 in the Jα helix with shorter Gly does not lead to any significant changes from the typical wild-type like behavior of the *To*AubZL_A1 variant.

*To*AubZL_A13 carries three mutations in the LOV domain – V450G, I483M, V499G. As mentioned earlier, the effect of the V499G mutation should not be as drastic as that of V499R. Val450 is largely conserved across most LOV photoreceptors, especially in all Aureos. However, in some cases, such as EL222 or PpSB1 LOV, it is replaced by threonine. The location of Val450 from Fα, in the vicinity of the flavin chromophore, is intriguing. The model structure reveals that Val450 actually refrains from directly interacting with the flavin moiety. Its non-polar side chain is approximately 4 Å away from the C5’ atom of the ribityl side chain. Even if replaced with threonine, as in the cases mentioned earlier, the CG2 atom would remain at an identical distance from C5’ atom of FMN, with no additional effect. Although the distance increases by 1.3 Å, substituting Val450 with the much shorter Gly neither significantly affects the structure of the LOV domain nor its photochemistry. Ile483 is positioned in close proximity to the hydrophobic part of flavin’s isoalloxazine ring. Therefore, the presence of non-polar amino acid is an absolute necessity for the proper incorporation of flavin. Indeed, this position is consistently occupied by a non-polar amino acid – most commonly leucine, valine, or isoleucine, and rarely methionine – in all LOV photoreceptors. Although the mutation of Ile483 to Met might have resulted in a steric clash with flavin, its flipping towards Fα instead made an additional hydrophobic contact with Ile457 while the Met483 CG atom maintained a favorable distance from the FMN (**Figure 3**). Interestingly, substitution by methionine i.e., L496M (Fuentes and Möglich, 2024) as well as the reverse i.e., M165I (Vaidya et al., 2011) have been extensively studied in *As*LOV2 and *Nc*VVD, respectively. The L496M variant of *As*LOV2 could be purified in soluble form with sufficient yield, and the rate constant value for dark recovery remained almost unchanged compared to wild-type *As*LOV2. This behavior is exactly the same as observed during the purification and dark recovery kinetics of the *To*AubZL_A13 variant. Similarly, the reverse M165I substitution in VVD resulted in a variant exhibiting dark recovery kinetics comparable to the wild-type. However, the doubly substituted M165I:M135I variant of VVD decelerated dark recovery by 10 orders of magnitude (Zoltowski et al., 2009). Our results, therefore, align extremely well with the rationally-designed mutants of *As*LOV2 and VVD. Among all the variants, *To*AubZL_A13 exhibited a slightly increased photocycle kinetics.

### 3.3. *In vitro* protein-DNA interaction study

After studying the photochemical properties, electrophoretic mobility shift assays (EMSA) were carried out to assess the DNA-binding affinity of the mutants, as efficient binding to a specific DNA substrate was crucial for the proper functioning of aureochrome and for its application as an optogenetic tool. EMSAs were performed with the purified and desalted samples of the *To*AubZL variants to investigate protein-DNA interaction patterns under both BL illumination and in the dark. The representative EMSA gel images are presented in (**Figure 4a, 4b)** and the DNA-binding profiles and calculated K_D_ values (**SI-5**) of all the variants are documented in **Table 2**. In all cases, a shift from no binding to substrate DNA-binding can be observed with increase in protein concentration. This is indicated by the increasing intensity of bands corresponding to free, unbound DNA as the protein concentration decreases. The complete absence or the appearance of very faint free DNA bands suggests protein-DNA complex formation, whereas prominent free DNA bands imply the inability of the protein to bind with its substrate DNA. A previous study in the lab (Deb et al., 2025) revealed that the wild-type *To*AubZL exhibited enhanced DNA-binding in the presence of blue light. All the mutated variants studied except *To*AubZL_A3, *To*AubZL_A12, and *To*AubZL_D1 (**Figure-4a**) showed a two-fold difference in DNA-binding affinity between light and dark conditions, similar to the wild-type *To*AubZL (Deb et al., 2025) (**Figure-4b**). We therefore investigated the significance of the mutations in the different *To*AubZL variants with respect to their impact on DNA-binding affinity.

**Figure 4:**
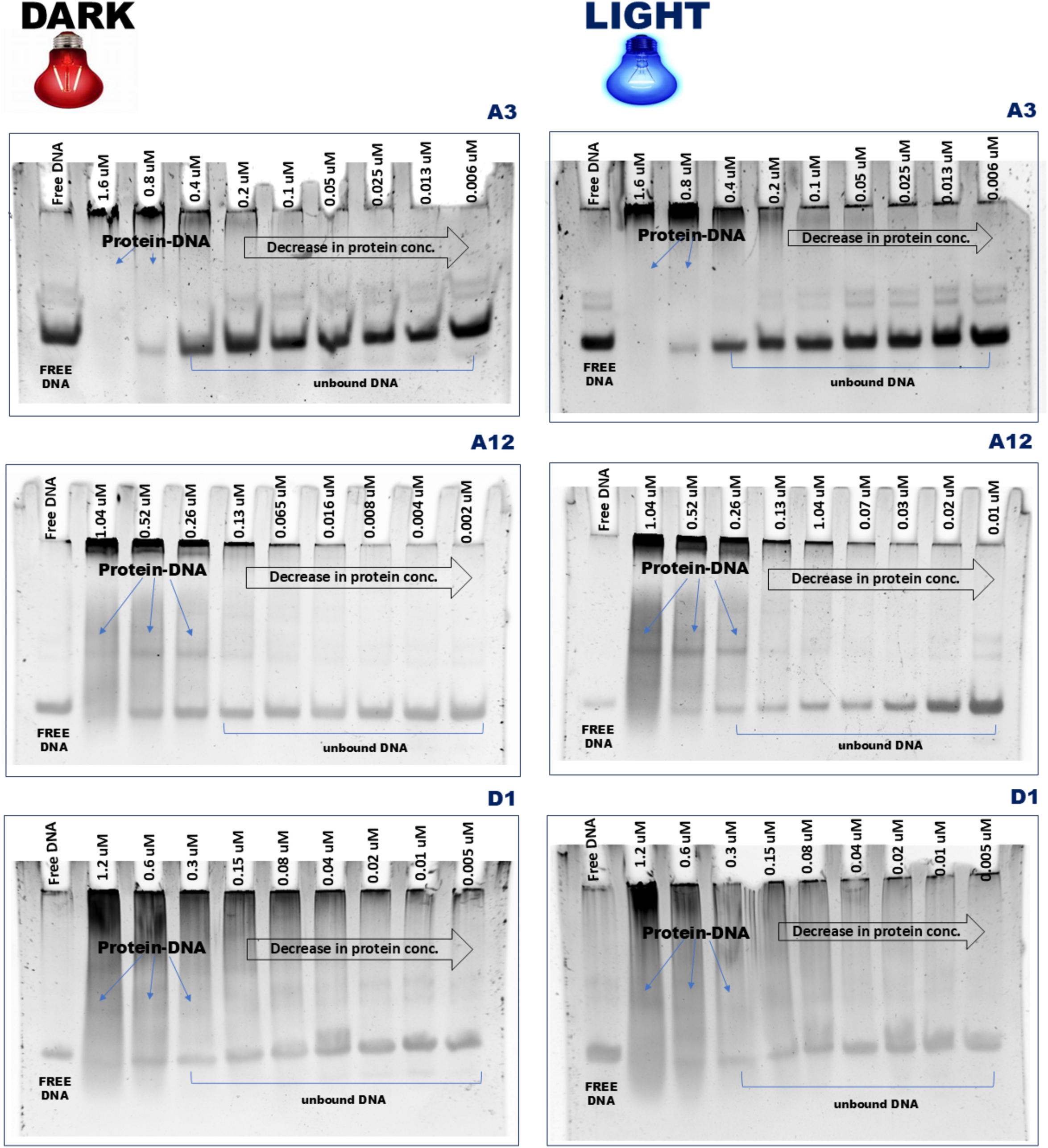
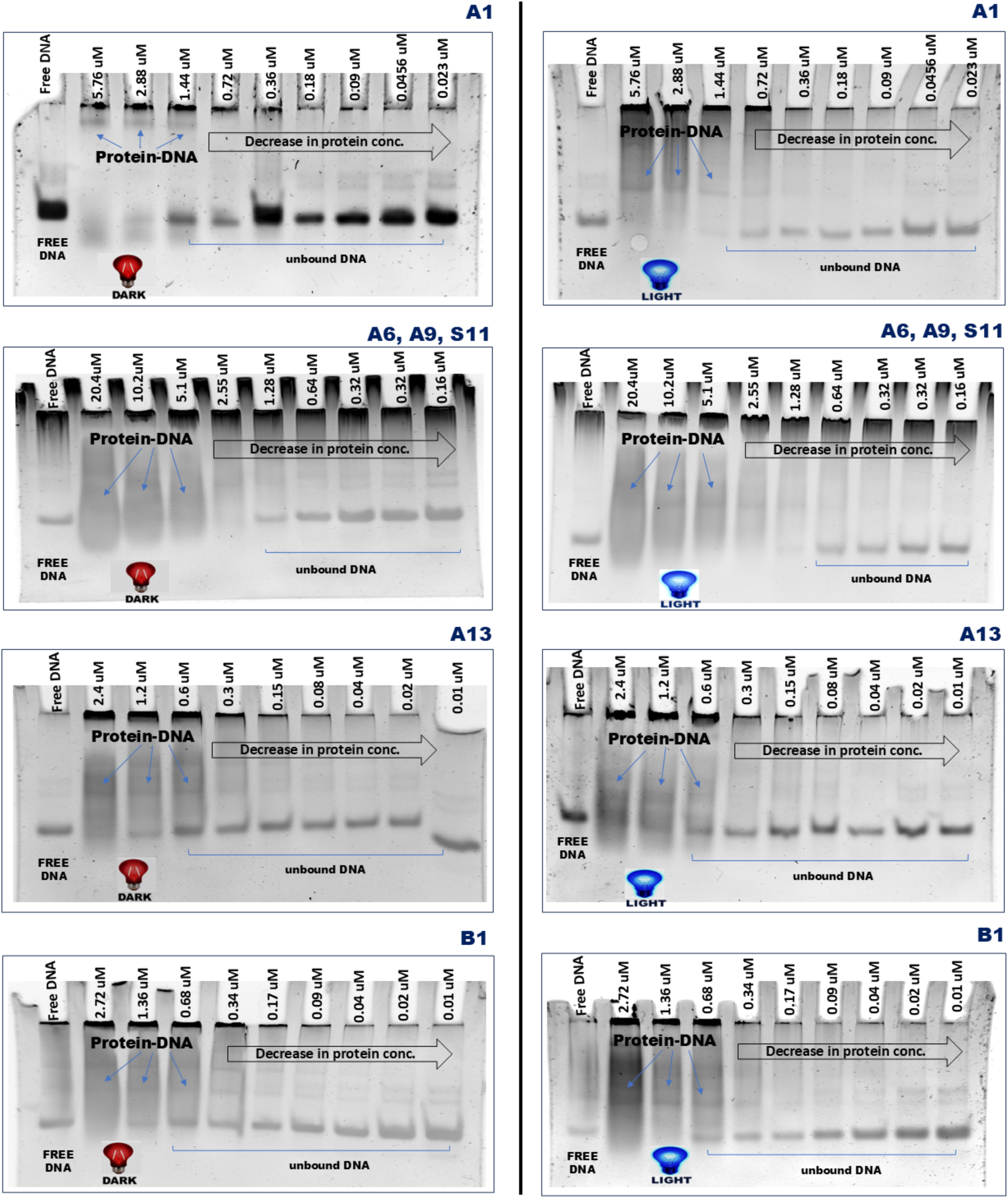
*In vitro* DNA-binding study of *To*AubZL variants by EMSA under light and dark conditions.

In the *To*AubZL_A1 variant, Lys318, which was primarily engaged with the phosphate backbone with the DNA substrate, is mutated to Gln. A mutation to another polar yet uncharged residue like glutamine (**Figure 5**), is not expected to alter the DNA-binding activity. As observed in the *To*Aureo1 docked structure (Deb et al., 2025), the mutation of Lys318 to glutamine retains polar contact with the phosphate backbone of the substrate DNA (DG’9).

**Figure 5:**
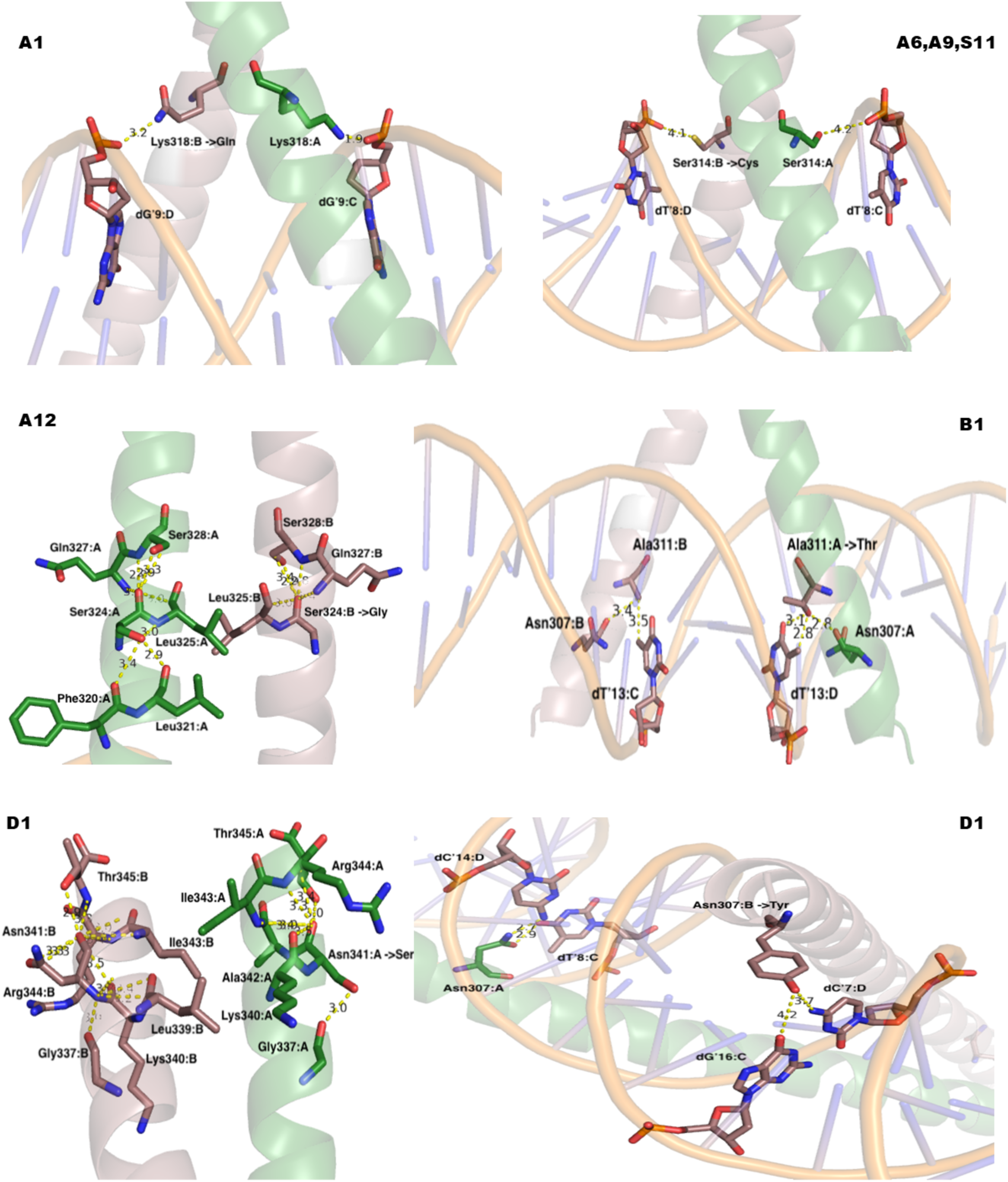
Structural details of the mutations introduced in the bZIP domain of the *To*AubZL variants.

The Ser314 residue, located in the signature basic region of aureochrome bZIP, was mutated to Cys in three variants – *To*AubZL_A6, *To*AubZL_A9, and *To*AubZL_S11. Barring Aureo2, this position is occupied either by Ser or Cys in all bZIPs. Therefore, serine-substituted basic region motifs are common. In the *To*Aureo1 S314C mutant, the native intra-helical hydrogen bonding interactions are preserved, without altering existing proximity with the DNA substrate (**Figure 5**), resulting in wild-type-like DNA-binding properties.

Ser324, a residue conserved in all Aureo1 homologs but not in other bZIPs, is mutated to Gly in *To*AubZL_A12. While this position is mostly occupied by polar residues, in some cases like PAP1 bZIP, an alanine is found. Located in a helical turn, a polar side chain would generally be more favourable at this position for interaction with the external solvent. However, substitution with the smallest amino acid, glycine, appears to have little consequence for two reasons – first, glycine is perfectly suited to reside in a helical turn (**Figure 5**); and second, with no Cβ, glycine has nothing but its polar backbone for being solvent-exposed.

In *To*AubZL_A13, the non-conserved Lys298 is located beyond the basic region of bZIP, which is primarily responsible for binding the substrate DNA. Nevertheless, given its proximity to the DNA-binding basic region as well as solvent exposure, substitution of Lys298 with positively-charged arginine – as seen in *To*AubZL_A13 – is not expected to hamper protein stability or interaction with the negatively-charged DNA.

The Ala311 residue is highly conserved in most bZIPs including Aureo1 homologs. However, in the bZIP of PAP1, it was replaced by glutamine similar to the Ala-to-Thr polar substitution found in the *To*AubZL_B1 variant. Ala311 is located adjacent to His310, which is a hallmark of all Aureos (Khamaru et al., 2025), with the exception of Aureo2 homologs. As mentioned by Khamaru et al., 2025, His310 is specifically involved in interaction with the substrate DNA, although it does not influence light-regulated DNA-binding. While the placement of Ala311 in the helical turn, especially between His and Lys, seems ideal; substitution with threonine appears to have no detrimental effects. As observed in the docked structure, the side chain of A311T (OG1) makes perfect contact with the nitrogenous base DT’13 (O4), thereby enhancing the specificity of the Aureo1–substrate DNA interaction (**Figure 5**).

*To*AubZL_D1 harbours two mutations in the bZIP domain – N307Y and N341S. The mutation of Asn307 is particularly interesting, as asparagine is absolutely conserved at this position across all bZIPs without any exception. Although the lack of difference in DNA-binding efficiency under light and dark conditions is perfectly explainable owing to its non-functional LOV domain, the unaltered DNA-binding affinity of *To*AubZL_D1 is quite surprising. Careful examination of the docked structure reveals that Asn307 formed a precise base-specific contact with DC’14 (N4) as well as DT’8 (O4) from the complementary strands of the DNA substrate (**Figure 5**). Upon mutation to Tyr, these particular interactions are lost. However, N307Y establishes a new base-specific contact with DC’7 (N4) and DG’16 (O6) from the complementary DNA strands. Very similar to our finding, mutation of the homologous Asn235 to Trp in GCN4 resulted in a mutant that behaved indistinguishably from the wild-type bZIP (Tzamarias et al., 1992) for a specific DNA substrate. However, for different promoter sequences, especially *in vivo*, mutations at the invariant Asn307 position could have significant impact. Asn341, in contrast, is poorly conserved among bZIPs and is located in the zipper region responsible for dimerization. Although buried Asn residues in the zipper region can be significant, the side chain of Asn341 is exposed to the external solvent. This further participates in intra-helical hydrogen bonding, and does not contribute to the dimer interface. Given its solvent exposure and retention of intra-helical hydrogen bonding interaction, albeit with different residues, the substitution of Asn341 with another polar residue, serine, is likely inconsequential.

Mutations in the DNA-binding motif and dimerization domain of the effector bZIP show little impact on the DNA-binding efficiency. The comparison of the BL-regulated DNA-binding affinity of the mutants based on the K_D_ value **Table 3** revealed that the variants *To*AubZL_A1 and *To*AubZL_A13 showed a maximum three-fold difference in DNA-binding efficiency between light and dark conditions.

**Table 3:**
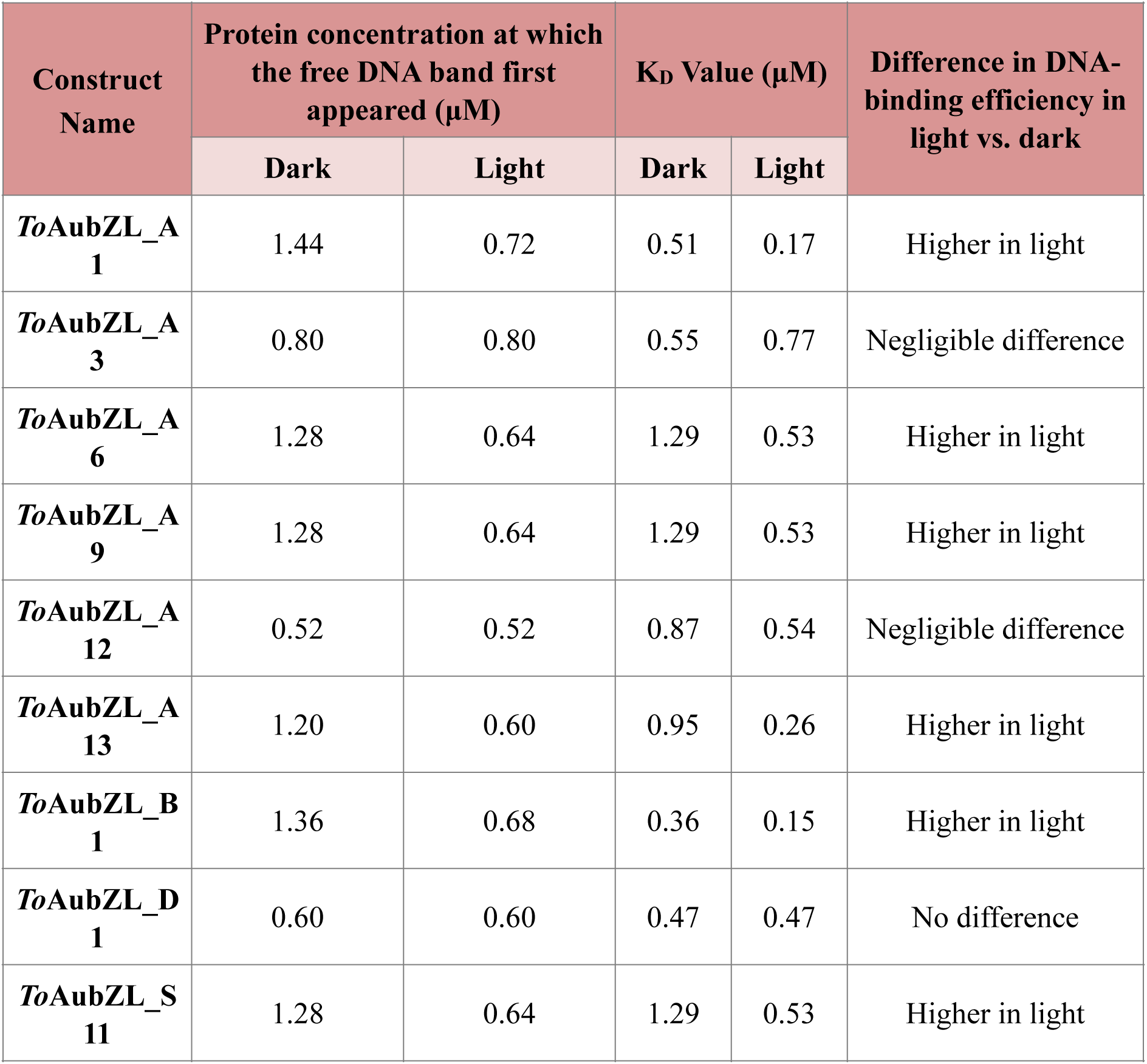
DNA binding affinities (K_D_ value) of the different *To*AubZL variants.

| Construct Name | Protein concentration at which the free DNA band first appeared ($\mu\text{M}$) | | $K_D$ Value ($\mu\text{M}$) | | Difference in DNA-binding efficiency in light vs. dark |
| --- | --- | --- | --- | --- | --- |
|  | Dark | Light | Dark | Light |  |
| *ToAubZL_A*<br>**1** | 1.44 | 0.72 | 0.51 | 0.17 | Higher in light |
| *ToAubZL_A*<br>**3** | 0.80 | 0.80 | 0.55 | 0.77 | Negligible difference |
| *ToAubZL_A*<br>**6** | 1.28 | 0.64 | 1.29 | 0.53 | Higher in light |
| *ToAubZL_A*<br>**9** | 1.28 | 0.64 | 1.29 | 0.53 | Higher in light |
| *ToAubZL_A*<br>**12** | 0.52 | 0.52 | 0.87 | 0.54 | Negligible difference |
| *ToAubZL_A*<br>**13** | 1.20 | 0.60 | 0.95 | 0.26 | Higher in light |
| *ToAubZL_B*<br>**1** | 1.36 | 0.68 | 0.36 | 0.15 | Higher in light |
| *ToAubZL_D*<br>**1** | 0.60 | 0.60 | 0.47 | 0.47 | No difference |
| *ToAubZL_S*<br>**11** | 1.28 | 0.64 | 1.29 | 0.53 | Higher in light |

## Discussion

Identification of the appropriate target for mutagenesis remains a crucial prerequisite for design and engineering proteins with desired functions. And literally, it is almost impossible to scan each and every amino acid residue of a protein (unlike the historic feat of introducing 2015 mutations in lysozyme and characterizing them!) to validate sequence-structure-function relations. However, an artificial intelligence based ‘protein inverse folding model’ has recently generated an artificial LOV sensor based on the LOV2 sequence from *Arabidopsis* phototropin. Even with a meagre 51% identity with the template sequence, this artificial LOV exhibited perfect photochemistry while achieving higher thermal stability (Herzog et al., 2025) – re-emphasizing the wonder of ‘the protein folding problem’. Site directed mutagenesis is usually carried out on obvious amino acid positions, which are either supposedly evolutionarily conserved or experimental evidence is available in homologous proteins. It is therefore challenging to accurately predict the effect of a specific mutation in advance. Nonetheless, nature being the greatest chemist is able to give rise to novel proteins through accumulation of mutations as an inevitable evolutionary process. Hence, a directed evolutionary approach through the introduction of random mutations has been employed to mimic natural evolution (Dennett, 1995). As the existence of functional proteins drives evolution, mutagenesis is accompanied by screening of the sequences that retained the original function or acquired mild improvement in performance (Arnold, 2018). Most of the mutations are either neutral or with minute changes, but some are even deleterious. Nevertheless, consecutive rounds of random mutation and subsequent screening can accumulate advantageous mutations to achieve enhanced functionality (Babbitt and Gerlt, 2000). In this way, this process can be channeled into an evolutionary route that sustains functionality through continuous pruning of nonviable mutations. Directed evolution thus proves to be a promising method for the optimization of proteins to achieve functions that are not observed in nature (Chen and Arnold, 2020). However, the greatest challenge lies in discovering proteins with refined functionality within a brief time frame. Iterations of mutagenic cycles followed by screening of the superior variant may take years of effort. Majority of the natural proteins are the products of gradual changes accumulated over billions of years. In many cases, the resultant proteins after mutagenesis can exhibit a maximum alteration of only 1% (Arnold, 2019). Even most of these modified sequences are not at all useful due to failure in functional protein production. Therefore, it is natural that we did not get variants with enhanced efficacy after screening a single generation library, generated through multiple mutagenic conditions. As our study represents the result from single generation directed evolution of *To*AubZL protein, subsequent rounds of mutations on the variants like *To*AubZL_A1 and *To*AubZL_A13 followed by multiple iterations of this process could produce improved variants.

While generating variant proteins with improved functions in a shorter time-scale is the major prerogative of directed evolution, a single step random mutagenesis allows us to search for those amino acid positions, which one would never explore rationally. Such amino acid residues may not necessarily bring out improved functional traits but could be helpful to understand the significance of them in protein function. One such finding is Ile414 in *To*AubZL, which is found to be crucial for FMN binding but was not reported earlier. Therefore, the major objective of this study has been to understand the sequence-structure-function correlations by nudging the fitness landscape, despite the availability of multiple structure-guided rationale-driven protein engineering techniques.

This study further highlights the relationship between single nucleotide polymorphism (SNP) and evolution. We observe that in all the variants obtained in this study, the substitution of amino acids is the result of single nucleotide change. Single nucleotide change in DNA can profoundly influence protein function through alteration in protein expression, folding, stability and catalytic activity (Paul, 2025). In nature, SNP is the prevalent source of genetic variation. It is found to occur naturally in every 1000 nucleotides to contribute uniqueness among individuals (Pocrnic et al., 2024). In humans, SNPs provide phenotypic diversity that contribute to differences in disease susceptibility and drug response (Zhong et al., 2003). SNP happens as a result of environmental pressure and leads to genetic isolation and divergence among species. Thus, the occurrence of SNP upon random mutation induced by differential concentration of MgCl_2_ and MnCl_2_ validates our idea of mimicking evolution by epPCR for the generation of variants. This study therefore can be a stepping stone towards the production of a superior variant of *To*AubZL protein to be used in optogenetics.

## Author Contributions

**Madhurima Khamaru:** Data curation, Investigation, Methodology, Formal analysis, Validation, Visualization, Writing – original draft, Writing – review and editing; **Pracheta Pal:** Data curation, Investigation, Methodology, Formal analysis, Validation, Visualization, Writing – original draft, Writing – review and editing; **Saptarshi Pal:** Data curation, Investigation, Methodology, Validation, Visualization, Writing– original draft; **Devrani Mitra:** Conceptualization, Supervision, Funding acquisition, Project administration, Investigation, Methodology, Formal analysis, Validation, Visualization, Writing – original draft, Writing – review and editing.

## Acknowledgements

MK acknowledges University Grants Commission (UGC) for her doctoral research fellowships. DM acknowledges funding from DBT, Government of India (BT/PR26435/BRB/10/1627/2017) for supporting the research work during the initial phase and SERB, Govt of India (CRG/2023/006463) at the present phase. The authors acknowledge the central instrumentation facility as well as the computer facility of the department supported by DBT-BUILDER and DST-FIST programs respectively. The authors also thank Debarshi Bose, Esha Moitra, Soumyajit Sardar and Abhilasha Ghosh for their cooperation.

