## Supplementary Information for "Random Mutagenesis for the Generation of Repertoire of Aureochrome-based Optogenetic Scaffolds"

### SI- 1. Details of the 4 rounds of error-prone PCR and subsequent cloning into pET28a.

| | $MnCl_2$ and $MgCl_2$ Concentration used during Error-Prone PCR | No. of Colonies Screened by Colony PCR | No. of Constructs Containing Mutation(s) in bZIP, Linker, and LOV Domains |
|---|---|---|---|
| **Error-Prone PCR Round 1** | 0.1 mM $MnCl_2$ | 14 (A1–A14) | 6 (A1, A3, A6, A9, A12, A13) |
| | 0.1 mM $MnCl_2$<br>5 mM $MgCl_2$ | Not cloned (PCR product yielded very faint band) | |
| | 0.3 mM $MnCl_2$ | 2 (B1, B2) | 1 (B1) |
| | 0.3 mM $MnCl_2$<br>5 mM $MgCl_2$ | Not cloned (PCR product yielded very faint band) | |
| **Error-Prone PCR Round 2** | 0.1 mM $MnCl_2$ | 4 (P1–P4) | 0 |
| | 0.1 mM $MnCl_2$<br>3 mM $MgCl_2$ | No colony formed after transformation of ligation products | |
| **Error-Prone PCR Round 3** | 0.1 mM $MnCl_2$ | 10 (C1–C10) | 0 |
| | 0.3 mM $MnCl_2$ | 10 (D1–D10) | 1 (D1) |
| | 0.1 mM $MnCl_2$<br>2 mM $MgCl_2$ | 8 (R1–R8) | 0 |
| **Error-Prone PCR Round 4** | 0.1 mM $MnCl_2$ | 8 (M1–M8) | 0 |
| | 0.3 mM $MnCl_2$ | 5 (N1–N5) | 0 |
| | 0.1 mM $MnCl_2$<br>2 mM $MgCl_2$ | 14 (S1–S14) | 1 (S11) |

**SI-2. Phylogenetic tree of LOV domain of all the *To*AubZL variants with other LOV sensors**

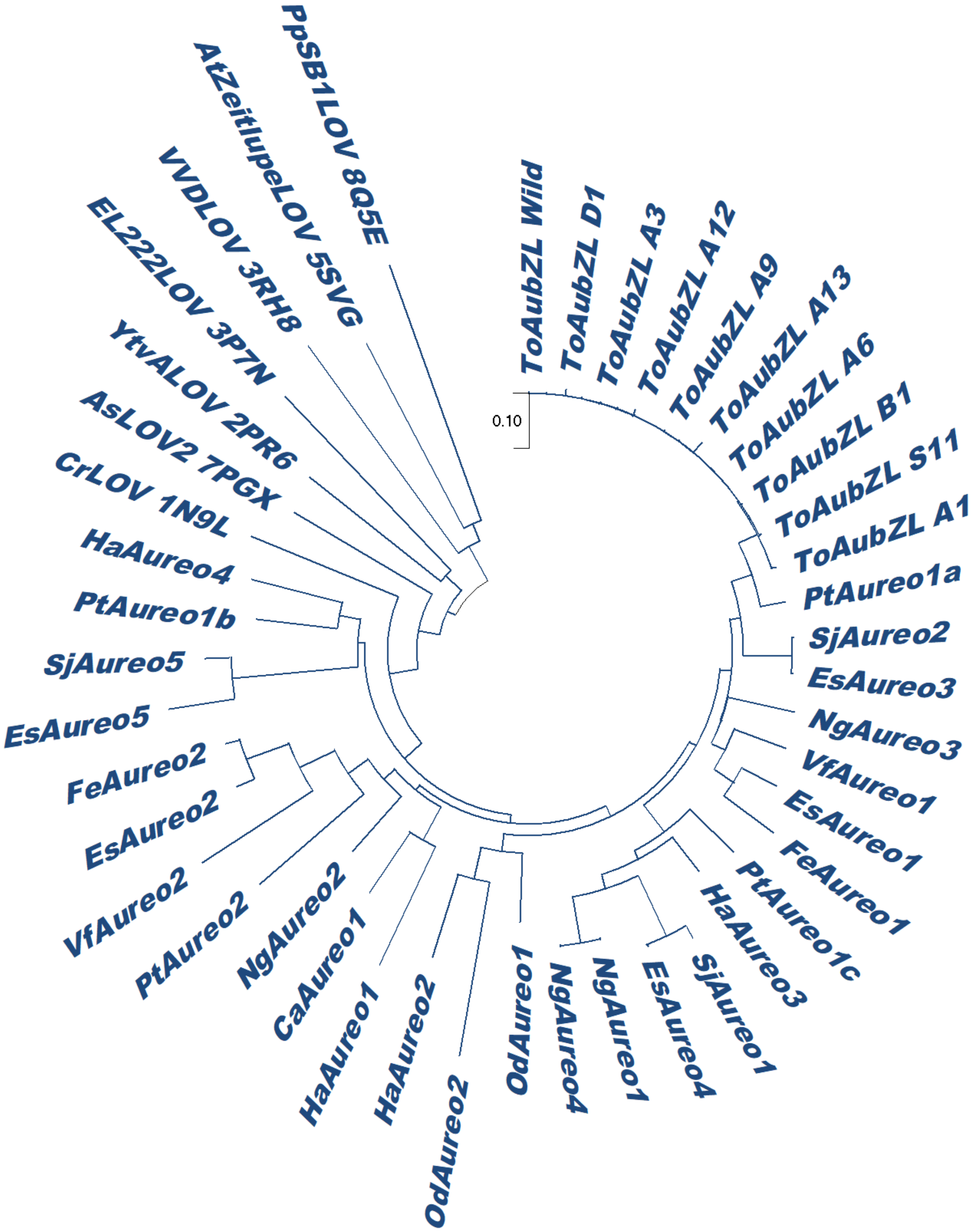

**SI-3. Phylogenetic tree of bZIP domain of all the *To*AubZL variants with other bZIPs**

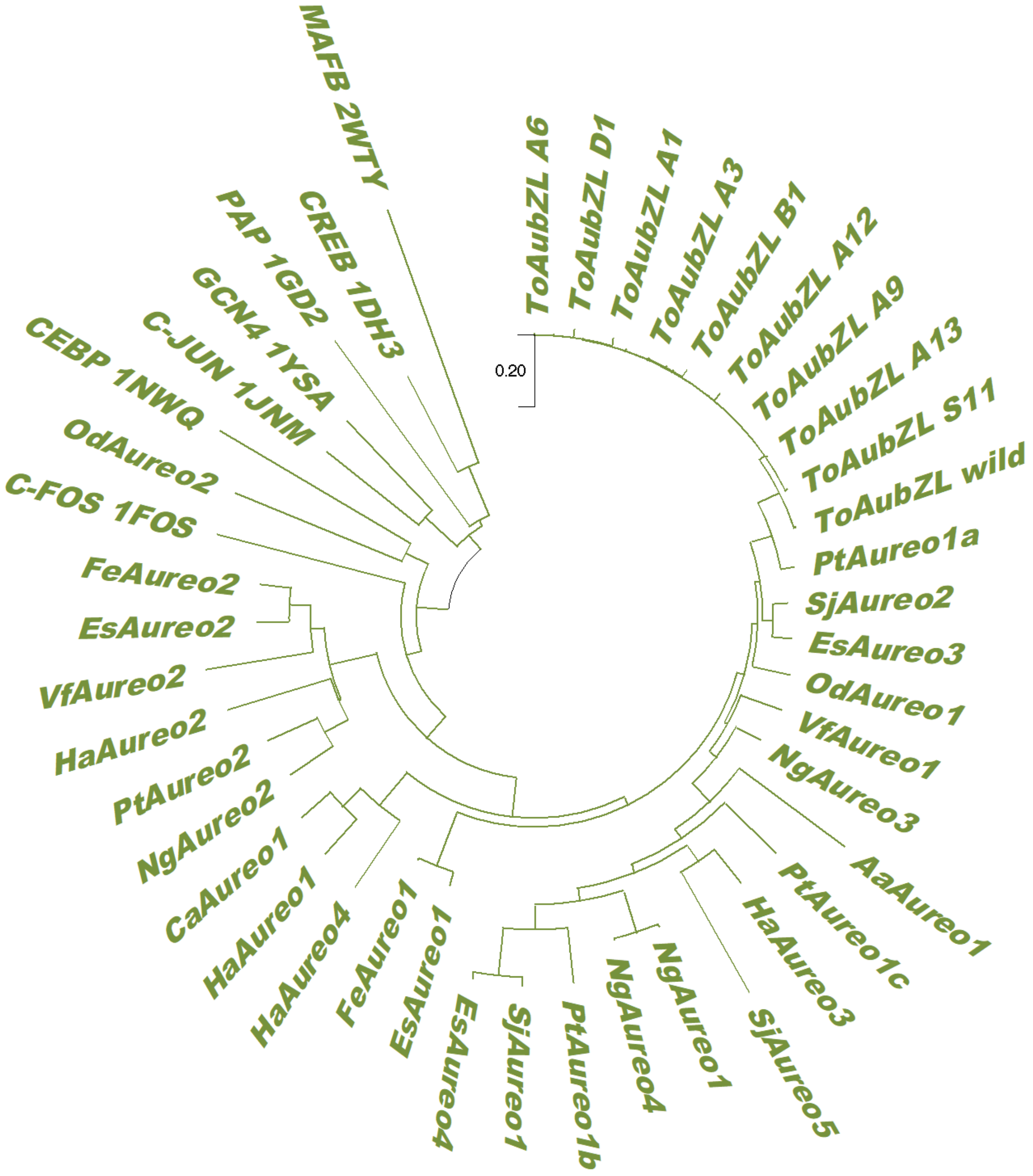

**SI-4. Rate constant calculation of the photocycle kinetics of the *To*AubZL variants**

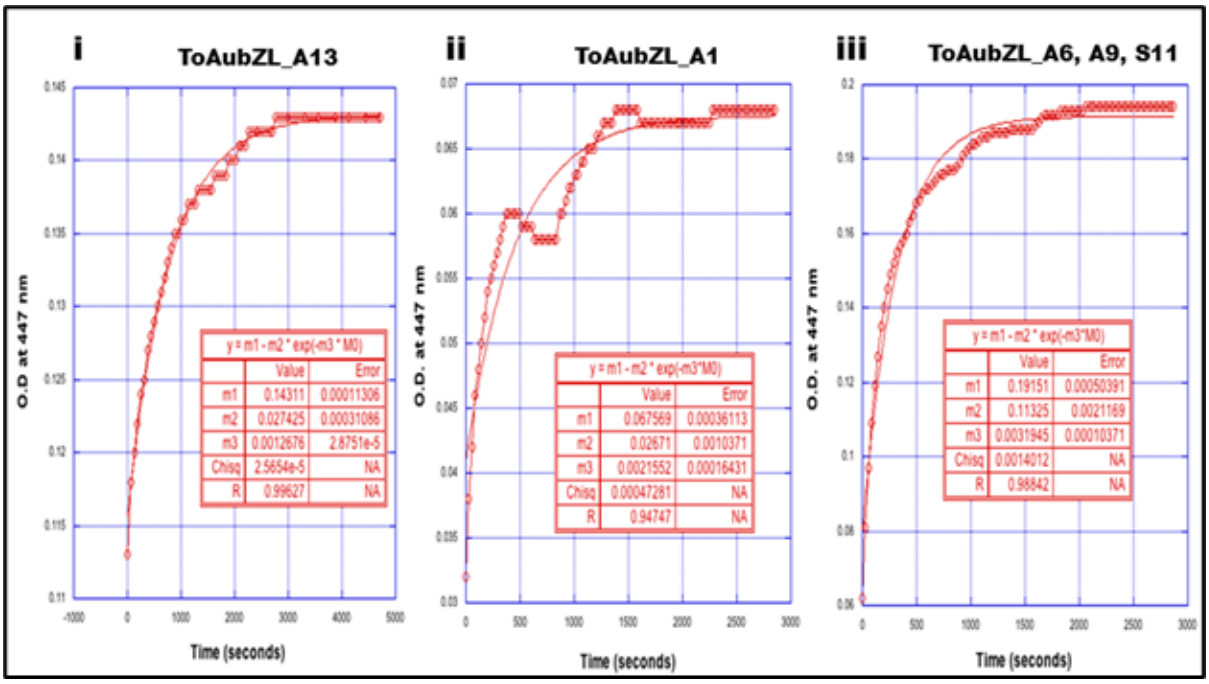

### SI-5. $K_D$ value calculation of the DNA binding affinity of *To*AubZL variants

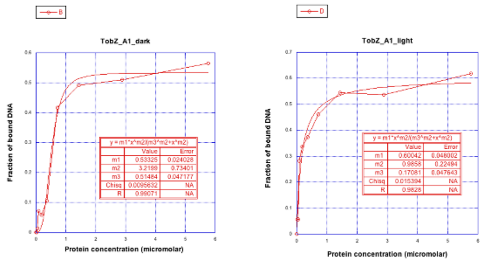


*To*AubZL_A1

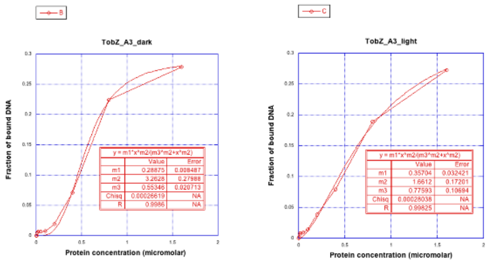


*To*AubZL_A3

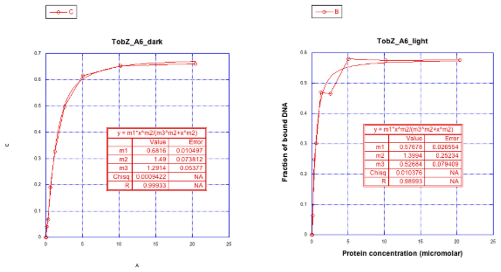


*To*AubZL_A6

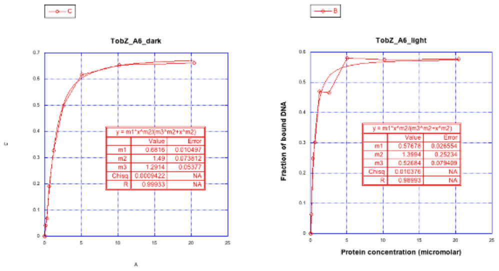


*To*AubZL_A9

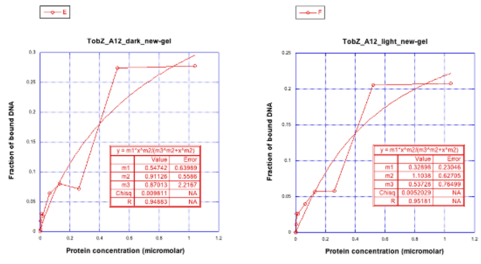


*To*AubZL_A12

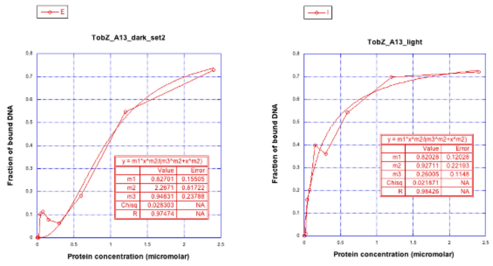


*To*AubZL_A13

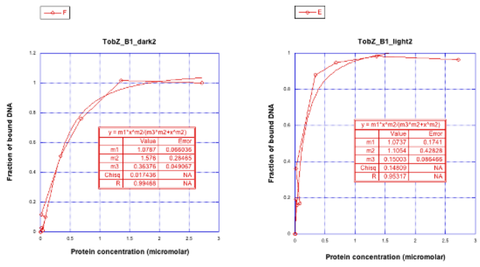


*To*AubZL_B1

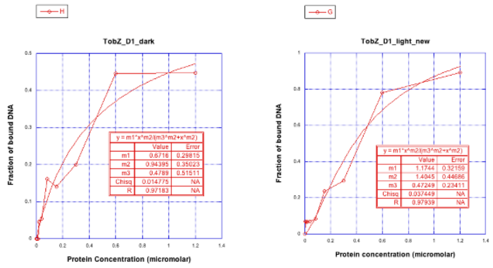


*To*AubZL_D1

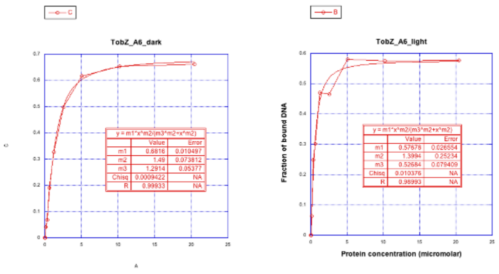


*To*AubZL_S11
